# Carrierwave: A granular, incentive-aligned infrastructure for scientific communication

**DOI:** 10.64898/2026.03.01.708795

**Authors:** Ido Bachelet

## Abstract

The peer-reviewed journal article imposes structural constraints on the dissemination, validation, and reuse of research outputs. Intermediate results, negative findings, methodological refinements, and replication attempts are systematically underrepresented in published literature, limiting visibility into ongoing research activity for both scientists and mission-driven funders. Here we present Carrierwave, an open infrastructure for continuous, granular scientific communication built on structured research objects (ROs), cryptographic provenance, blockchain-based attribution, and programmable incentive mechanisms. Each RO represents an atomic unit of scientific output -- a single experimental result, negative finding, dataset, protocol, or replication -- that is hashed for content integrity, stored in a persistent database, and optionally minted as an ERC-721 non-fungible token on the Ethereum blockchain. The system includes an on-chain bounty pool enabling funders to directly incentivize specific research activities, and an automated analysis layer that synthesizes disclosed ROs into continuously updated research landscape maps. We describe the system architecture, report on its implementation and deployment on Ethereum mainnet, and present a quantitative analysis of disease-specific publication frequency demonstrating the information latency problem that Carrierwave addresses. The distribution of publication frequency across disease areas is highly skewed, with the majority of conditions represented by fewer than four publications per year in high-impact biology journals. For diseases in the long tail, the interval between successive publications may span months or years. Publication frequency correlates poorly with disease burden, instead reflecting historical research community size and advocacy momentum. By reducing the unit of communication to the individual research object and eliminating editorial gatekeeping as a prerequisite for disclosure, Carrierwave increases the effective sampling rate of scientific activity in precisely the domains where publication-based visibility is most sparse. The system is live at https://carrierwave.org.

## Introduction

The peer-reviewed journal article has served as the primary vehicle for scientific communication since the seventeenth century. Despite the proliferation of preprint servers, open-access mandates, and digital repositories, the journal article remains the dominant unit of scientific discourse, career evaluation, and funding allocation [1,2]. This system, while historically productive, imposes structural constraints that are increasingly recognized as impediments to scientific progress. The journal article bundles multiple experimental results into a narrative package optimized for editorial acceptance rather than for maximal transparency or reuse. The peer review and editorial process introduces substantial temporal delays between the completion of experimental work and its public availability [3,4]. The incentive structure of journal publishing systematically favors novel, positive findings over negative results, replications, and incremental methodological contributions [5,6]. The resulting “file-drawer problem” [7] and the broader reproducibility crisis [8,9] are widely acknowledged consequences of these structural incentives.

Large-scale replication efforts have quantified the magnitude of this problem. The Open Science Collaboration’s attempt to replicate 100 psychology studies found that only 36% produced statistically significant results consistent with the original reports [9]. In preclinical cancer biology, the Reproducibility Project reported that effect sizes in replications were on average 85% smaller than those in original studies [10]. The file-drawer problem is structural rather than incidental: the incentive architecture of journal publishing rewards novelty and statistical significance while offering little reward for negative results, null findings, or exact replications [6,11].

These limitations affect not only researchers but also mission-driven funders -- disease foundations, patient advocacy organizations, and public funding agencies -- whose strategic decisions depend on timely, comprehensive visibility into the research landscape. For diseases with small research communities or low publication volume, the effective sampling rate of published literature may be measured in years rather than months, rendering entire domains of active research effectively invisible to the organizations funding them.

Several reform efforts have sought to address aspects of this problem. Preprint servers such as bioRxiv have reduced time-to-disclosure for completed manuscripts [12]. Open-access mandates have improved accessibility [13]. Micropublication journals have demonstrated the value of publishing granular results including negative findings [14]. The FAIR data principles have articulated standards for findable, accessible, interoperable, and reusable research outputs [15]. Decentralized science (DeSci) initiatives have explored blockchain technology for research attribution and funding [16,17]. Nanopublications have formalized individual scientific assertions as machine-readable, attributed statements [18,19], and blockchain-based proof-of-existence has been discussed for establishing scientific priority [20]. Prize-based incentive models have been formalized in the context of innovation inducement prizes [21], and registered reports have attempted to decouple publication incentives from outcomes [11,22].

However, none of these efforts address the full set of constraints simultaneously. Preprint servers accelerate disclosure but do not change the unit of communication. Micropublication reduces article size but retains editorial gatekeeping and lacks on-chain provenance or economic incentive layers. FAIR principles describe desirable properties of data but do not specify an integrated system for disclosure, validation, and incentive alignment. DeSci projects have explored tokenized funding and IP-NFTs but have generally not addressed the granularity of scientific output or the specific information needs of mission-driven funders.

This paper presents Carrierwave, an integrated infrastructure that addresses these limitations through five coordinated design choices: (i) structured research objects as the atomic unit of scientific communication; (ii) cryptographic hashing and blockchain-based provenance for priority and attribution; (iii) an explicit relationship graph among research objects enabling continuous validation tracking; (iv) a programmable bounty pool that aligns funder incentives with specific research activities; and (v) an AI-powered analysis layer that synthesizes disclosed research objects into navigable research landscape maps. The system is implemented as a web application deployed on Ethereum mainnet and is publicly accessible at https://carrierwave.org.

## Materials and methods

To quantify the information latency problem motivating this work, we analyzed disease-specific publication frequency across high-impact biology and biochemistry journals. Journal lists comprising 200 journals each were obtained from the Observatory of International Research (OOIR) Biology & Biochemistry rankings, with journals ranked by number of papers and by impact factor. Article metadata (titles, publication dates, and MeSH terms) were retrieved from the PubMed database via the E-utilities API (esearch and efetch) for the preceding five years, with requests spaced by >=0.4 s and fetches performed in batches of up to 200 article IDs. Disease relevance was determined by matching article titles and MeSH terms against a curated set of approximately 150 disease terms (e.g., cancer subtypes, infectious diseases, neurological conditions) and by searching titles for 100 predefined rare diseases (e.g., Huntington disease, cystic fibrosis, Tay-Sachs disease). Articles without any disease match were excluded. For each disease, the number of matching articles was divided by the number of days in the analysis window (~1,826 days) to obtain publication frequency in papers per day, and diseases were ranked by this metric. Separate analyses were conducted for the main disease set and the rare disease set; results were combined for visualization. To assess whether publication frequency reflects disease burden, publication frequency was plotted against estimated global patient count for rare diseases.

To quantify time-to-disclosure under existing editorial models, we measured the interval between manuscript receipt and publication for recent articles across five biology journals spanning a range of editorial models: eLife (n=61), PeerJ (n=1,000), Current Biology (n=641), Scientific Reports (n=979), and MicroPublication Biology (n=989).

The Carrierwave system was implemented as a full-stack web application using Next.js with TypeScript and the App Router. Authentication uses Sign-In with Ethereum (SIWE) with iron-session for cookie-based session management. Research objects are stored in a key-value database (Vercel KV, backed by Redis) with associated metadata; uploaded figures and data files are stored in cloud blob storage (Vercel Blob). The on-chain layer consists of three smart contracts written in Solidity 0.8.28 using the OpenZeppelin library, compiled with the Hardhat 2 framework with optimizer enabled (200 runs). The interactive relationship graph uses D3.js for force-directed layout. The automated analysis layer uses the Claude language model (Anthropic) for landscape synthesis. The application is deployed on Vercel with continuous deployment from GitHub.

## Results

The distribution of disease-specific publication frequency is highly skewed (Fig 1). A small number of disease categories -- including tumor biology, Alzheimer’s disease, metastasis, and HIV -- dominate journal output at rates exceeding 1 paper per day. The distribution exhibits a long tail, with the majority of disease terms appearing at rates below 0.01 papers per day, corresponding to fewer than 4 publications per year. For diseases in the long tail, the interval between successive publications may span months or years. During these intervals, ongoing experimental work -- including negative results, feasibility studies, and methodological refinements -- remains invisible to the broader research community and to the funders supporting this work.

**Fig 1.**
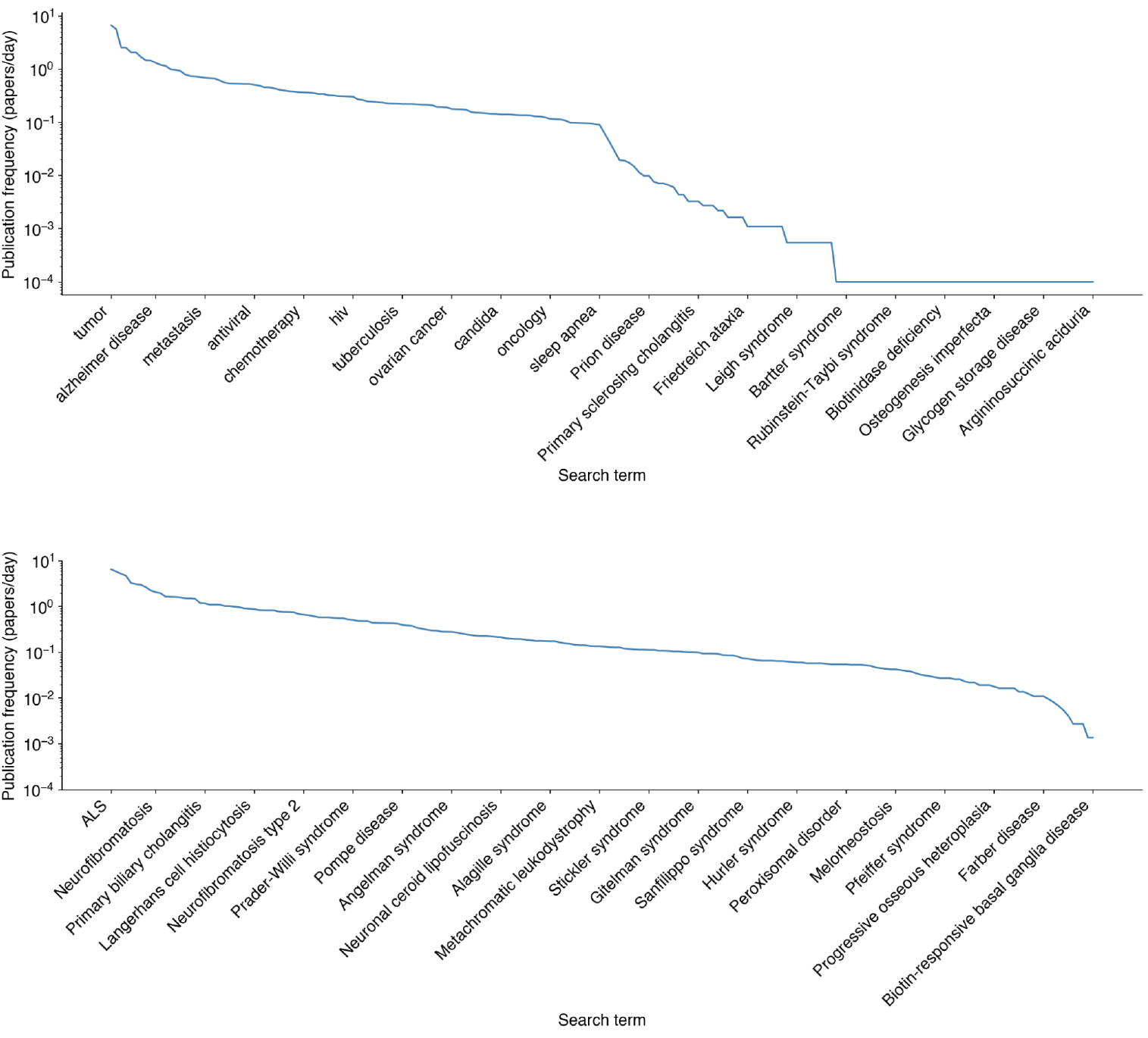
Disease-specific publication frequency in high-impact biology and biochemistry journals. Publication frequency (papers/day) over a five-year from 100 top ranked journals (top) or two-year across the entire PubMed (bottom) window for 200 disease-related search terms, displayed on a logarithmic scale. The distribution is highly skewed, with a small number of disease categories dominating journal output and a long tail of diseases appearing only sporadically.

The relationship between global disease prevalence and publication frequency is weak (Fig 2), with publication rates spanning two to three orders of magnitude at any given prevalence level. Diseases with strong advocacy infrastructure and established research communities -- notably cystic fibrosis and ALS -- achieve publication rates comparable to or exceeding those of diseases with 10- to 100-fold larger patient populations. Conversely, several high-prevalence conditions disproportionately affecting populations in lower-income countries, such as thalassemia and sickle cell disease, show publication rates below what their disease burden would predict. This disconnect demonstrates that journal publication frequency encodes historical research community size, advocacy momentum, and funding history rather than patient need.

**Fig 2.**
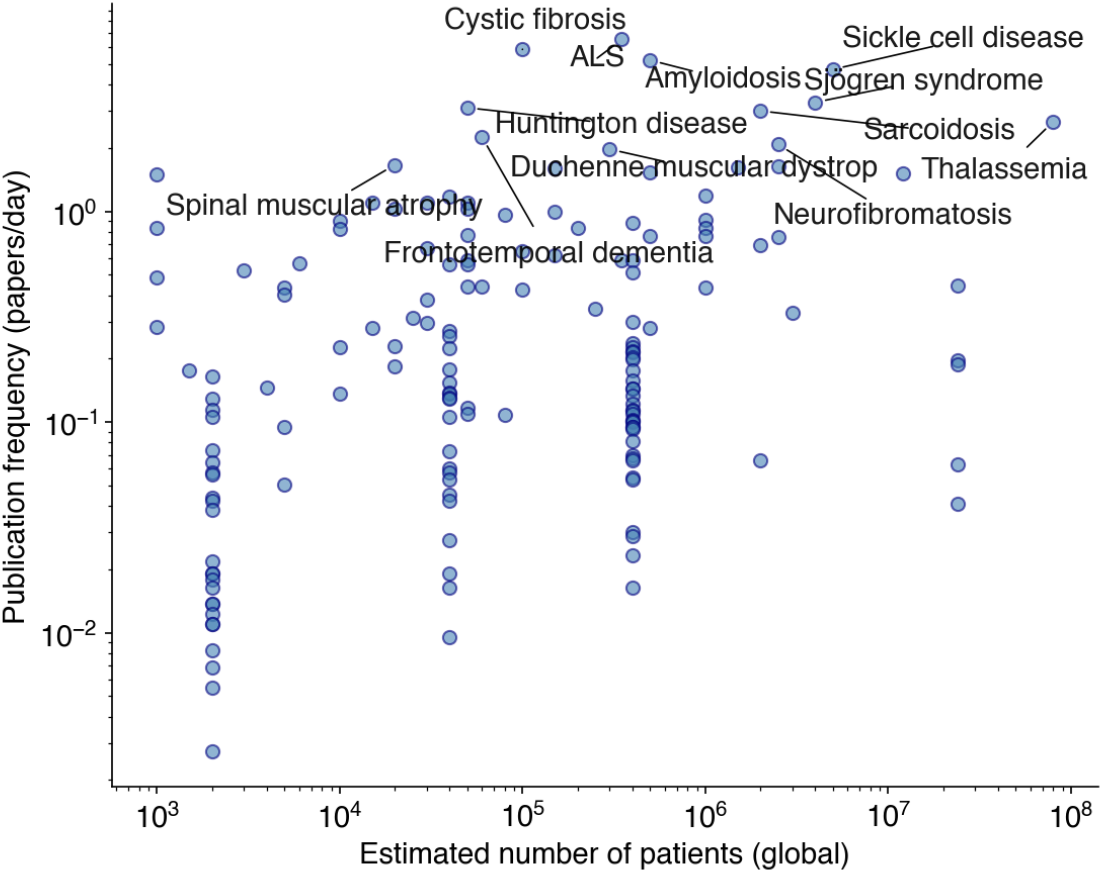
Relationship between global disease prevalence and publication frequency for rare diseases. Scatter plot with estimated global patient count (x-axis, log scale) against publication frequency (y-axis, log scale, papers/day). The diffuse scatter demonstrates that publication output is a poor proxy for disease burden.

Across the five journals examined, median submission-to-publication times fall in the range of approximately 50 to 300 days (Fig 3). Even journals designed for faster turnaround -- PeerJ and Scientific Reports -- show medians exceeding five months. MicroPublication Biology, which accepts granular results and is the closest existing analog to Carrierwave in terms of output size, achieves a lower median but retains a long tail extending past 500 days. Every journal exhibits substantial variance, with individual papers taking over 1,000 days from submission to publication.

**Fig 3.**
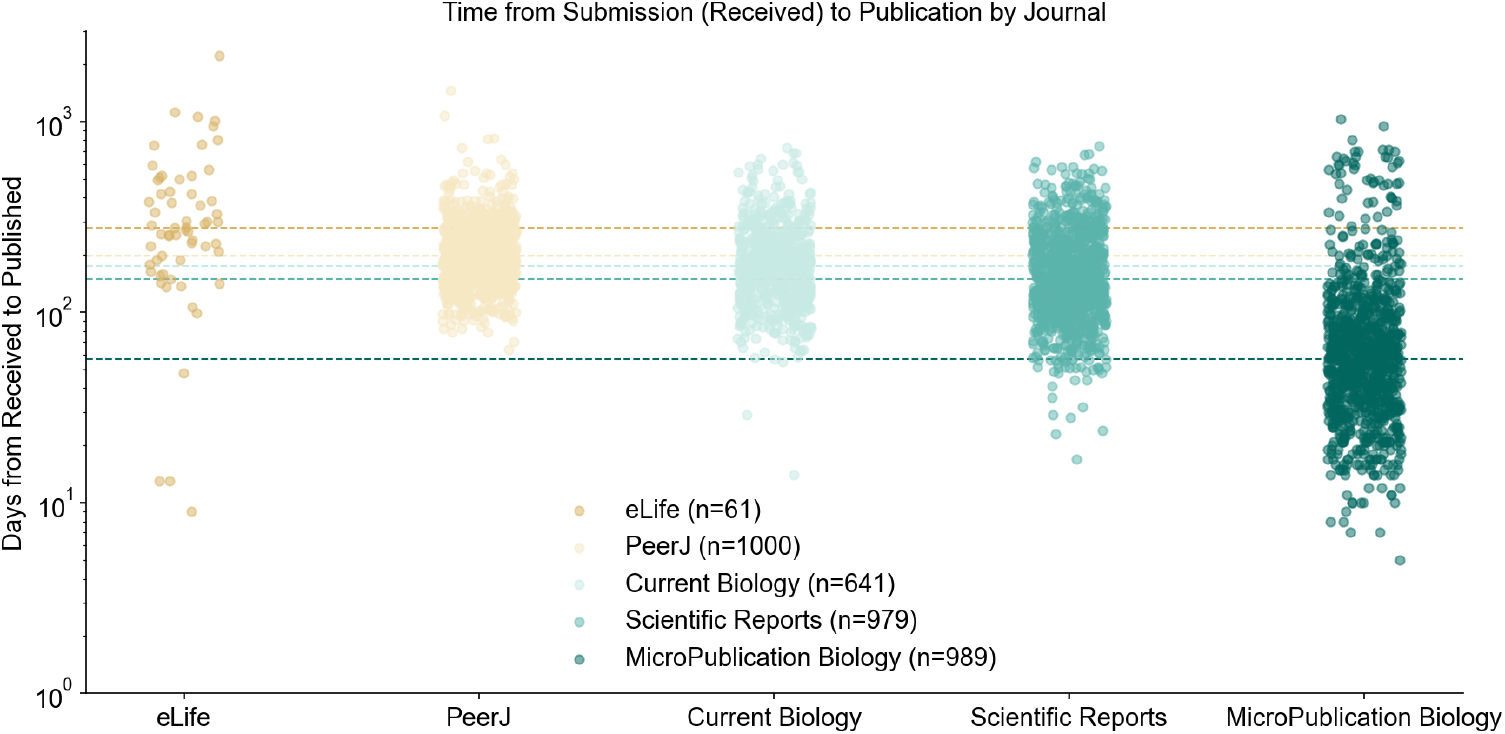
Time-to-visibility for scientific results under selected editorial models. Days from manuscript receipt to publication for recent articles in five biology journals: eLife (n=61), PeerJ (n=1,000), Current Biology (n=641), Scientific Reports (n=979), and MicroPublication Biology (n=989). Dashed horizontal lines indicate per-journal medians. Even journals designed for faster turnaround show medians exceeding five months, establishing a structural floor on information latency inherent to the journal system.

The Carrierwave system comprises five layers. The submission layer captures structured metadata through an eight-step guided wizard, including title, abstract, research object type, disease area tags, species, confidence level, figure upload, and relationship links to existing ROs. At submission, the system computes a SHA-256 hash of the RO content, and the submitting researcher authenticates via SIWE, binding the submission to a specific blockchain wallet address. The storage layer assigns each RO a universally unique identifier and indexes it by recency, submitter wallet, and disease area tags. A relationship graph encoding typed edges (replication, contradiction, extension, derivation, methodological reuse) between RO identifiers is maintained.

The on-chain layer consists of three smart contracts deployed on Ethereum mainnet and verified on Etherscan (Table 1). The CarrierwaveROv2 contract enables researchers to mint submitted ROs as ERC-721 non-fungible tokens, recording the RO identifier and content hash on-chain. A minting fee is forwarded to the CWTreasury contract, which implements a configurable distribution function splitting accumulated fees among designated recipients. The CWBountyPool contract implements the full lifecycle of funded research bounties: funders create bounties by specifying a disease area, evaluation criteria, and deadline while locking ETH in escrow; scientists submit claims by linking research objects; funders approve or reject claims and assign reward shares; upon finalization, the contract distributes funds to approved claimants with a 2.5% platform fee deducted to the treasury. An automatic institutional split mechanism accommodates researchers required to share revenue with employing institutions, with institutional shares held in escrow when no institutional wallet is registered.

**Table 1.** Deployed smart contract addresses (Ethereum mainnet).

| Contract | Address |
| --- | --- |
| Contract | Address |
| CWTreasury | 0xB93A8033883f7d9050bbfabEf35bD6a7D09d834a |
| CarrierwaveROv2 | 0xEe7c58E02387548f7628e467d862483Ebb285e7f |
| CWBountyPool | 0x63d28D915Ab0dF57a86547562d5f4A086d3b2A81 |

The discovery layer provides three interfaces: a paginated, filterable explorer feed; individual RO detail pages displaying full metadata, figures, content hash, minting status, and relationships; and an interactive force-directed graph visualization rendering all ROs as nodes and their typed relationships as directed edges. The analysis layer periodically scans all disclosed ROs and their relationships using a large language model to cluster ROs by disease area, identify areas of active investigation or stagnation, surface contradictory findings, suggest replication targets, and assess confidence trajectories. The complete technology stack is summarized in Table 2.

**Table 2.** Technology stack.

| Layer | Technology | Notes |
| --- | --- | --- |
| Layer | Technology | Notes |
| Web framework | Next.js (App Router) | TypeScript, server-side rendering |
| Authentication | Sign-In with Ethereum | SIWE protocol + iron-session |
| Database | Vercel KV (Redis) | Key-value storage with indices |
| File storage | Vercel Blob | Figures, data files |
| Blockchain | Ethereum mainnet | Chain ID: 1 |
| Smart contracts | Solidity 0.8.28 | OpenZeppelin, Hardhat 2 |
| Token standard | ERC-721 | Non-fungible provenance tokens |
| Graph visualization | D3.js | Force-directed layout |
| AI analysis | Claude (Anthropic) | Landscape synthesis |
| Deployment | Vercel | Continuous deployment from GitHub |

## Discussion

Carrierwave differs from existing approaches to scientific communication along several dimensions (Table 3). Traditional journal publishing and micropublication both rely on editorial gatekeeping prior to disclosure, produce implicit citation-based relationships, and offer retrospective funder visibility. Carrierwave replaces pre-publication gatekeeping with immediate disclosure followed by continuous post-hoc validation, provides an explicit typed relationship graph, and enables real-time funder visibility through an integrated economic layer.

**Table 3.** Comparison of Carrierwave with existing scientific communication approaches.

| Dimension | Traditional journal | Micropublication | Carrierwave |
| --- | --- | --- | --- |
| Unit size | Full paper | Small paper/result | Structured RO |
| Gatekeeping | Editorial + peer review<br>Peer review | Post first, continuous validation |  |
| Time to disclosure | Months to years | Weeks to months | Immediate |
| Provenance | DOI (post acceptance) | DOI | Cryptographic + blockchain |
| Relationships | Implicit citations | Implicit citations | Explicit typed graph |
| Validation model | One-shot peer review | One-shot peer review | Continuous multi-source |
| Incentive mechanism | Citations | Citations | Validations + reuse + bounties |
| Funder visibility | Retrospective | Retrospective | Real-time |
| Economic layer | None | None | Native (ETH) |

A defining feature of the system is the absence of pre-publication gatekeeping. Research objects are publicly visible immediately upon submission, with quality assessment occurring through post-hoc mechanisms: replication objects, confidence scoring, and reputation dynamics based on historical validation outcomes. This model inverts the traditional sequence of validation followed by publication. The primary benefit is the elimination of information latency: results become available for inspection, reuse, and replication without waiting for editorial approval. The primary risk is that unvalidated or low-quality submissions may be visible alongside well-supported findings. The system mitigates this risk through confidence level self-reporting at submission, continuous accumulation of validation evidence through the relationship graph, automated anomaly detection by the analysis layer, and an incentive structure that ties long-term reward to downstream validation rather than initial posting. This model is conceptually aligned with the “publish, then filter” paradigm discussed in the open science literature [23], extended with explicit economic incentives and structured validation tracking.

The use of Ethereum as a provenance layer provides immutability, public auditability, platform independence, and timestamping with blockchain-level security guarantees. On-chain provenance also introduces limitations: transaction costs may fluctuate, and on-chain storage is limited to hashes and identifiers, with underlying data remaining in off-chain storage. The environmental concerns associated with proof-of-work blockchains have been substantially mitigated by Ethereum’s transition to proof-of-stake, which reduced energy consumption by approximately 99.95% [24].

The on-chain bounty pool represents a mechanism for aligning funder and researcher incentives around specific scientific objectives. Unlike traditional grants, bounties specify concrete deliverables and release funds only upon delivery and funder approval. This design is influenced by the literature on innovation inducement prizes [21], adapted for the scientific research context. The key difference from prize models is the integration of bounties with a structured research object system: claims are linked to specific, inspectable ROs with provenance records. The automatic institutional split mechanism addresses a practical barrier to researcher participation by encoding revenue-sharing obligations at the smart contract level.

The publication frequency analysis presented here demonstrates that the information latency problem is particularly acute for diseases with small research communities -- precisely the domains where mission-driven funders are most active. Disease-focused foundations, patient advocacy organizations, and public-interest funding agencies operate under a structural information constraint: their visibility into the scientific process is largely mediated by peer-reviewed publications. Funders are often unable to observe unsuccessful or abandoned experimental approaches, incremental feasibility studies, early signals of convergence or contradiction across laboratories, and shifts in methodological practice prior to formal publication. Continuous granular disclosure transforms research oversight from a retrospective, publication-dependent process into near-real-time landscape analysis. The bounty pool enables funders to directly incentivize replications, negative result reporting, and methodological validation -- activities the journal system structurally undervalues. The automated analysis layer further supports decision-making by synthesizing disclosed research objects into field-level maps highlighting emerging trends, stalled lines of inquiry, and areas where independent validation may be particularly valuable.

Several limitations should be acknowledged. The system is currently scoped to non-human, non-sensitive research data; extension to clinical or human subjects data would require consent management, access control, and compliance mechanisms not present in the current architecture. The post-first, validate-later model depends on the emergence of a community of validators, and in the early stages of adoption the validation layer will be sparse. Institutional intellectual property constraints may limit what some researchers are able to disclose. The automated analysis layer relies on a large language model whose outputs are presented as interpretive aids, not as authoritative determinations of scientific validity. The current implementation relies on centralized infrastructure; migration toward decentralized storage is planned. Formal evaluation of the system’s impact on disclosure behavior, information latency, and research outcomes will require longitudinal study with a sufficient user base.

In summary, Carrierwave provides an open infrastructure for scientific communication that disaggregates the journal article into granular research objects, provides cryptographic and blockchain-based provenance, enables continuous validation through an explicit relationship graph, aligns funder and researcher incentives through a programmable bounty pool, and synthesizes disclosed outputs into continuously updated research landscape maps. The system is deployed on Ethereum mainnet and publicly accessible at https://carrierwave.org. It is designed to coexist with existing academic publishing pathways, addressing the disclosure of outputs below the journal publication threshold, the reduction of information latency, and the provision of real-time research visibility for mission-driven funders. By optimizing for signal density rather than narrative density, the system makes visible much of the research process currently hidden from view.

## Data availability

The Carrierwave platform is publicly accessible at https://carrierwave.org. Smart contract source code is verified and publicly readable on Etherscan at the addresses listed in Table 1. The publication frequency analysis code and data are available from the author upon request.

